# Disease Stage- and Risk-Associated RNA Editing Signatures in Acute Myeloid Leukemia and Their Utility for Peripheral Blood-Based Assessment

**DOI:** 10.64898/2026.06.29.735403

**Authors:** Tongjun Gu, Doan Bui, Ji-Hyun Lee

**Affiliations:** Versiti Blood Research Institute, Milwaukee, Wisconsin; Data Science Institute, Medical College of Wisconsin, Milwaukee, Wisconsin; Department of Biomedical Engineering, Medical College of Wisconsin and Marquette University, Milwaukee, Wisconsin; Department of Biostatistics, University of Florida, Gainesville, Florida; Cancer Quantitative Sciences, UF Health Cancer Institute, Gainesville, Florida

## Abstract

RNA editing is a widespread post-transcriptional regulatory mechanism, but its role in acute myeloid leukemia (AML) remains incompletely understood. We analyzed RNA editing in 59 paired diagnosis-relapse AML samples and eight age-matched healthy controls using a stringent discovery pipeline and beta-binomial regression framework accounting for overdispersion and repeated measurements. A total of 166,323 high-confidence RNA editing sites mapping to 5,917 genes were identified. Of tested sites, 1.2%–3.6% varied significantly by disease stage or ELN-2022 risk group. Disease stage-specific editing signatures distinguished healthy controls, diagnosis, and relapse samples, with relapse-associated signals validated in an independent AML cohort. ELN-2022 risk-specific editing signatures showed substantial overlap between intermediate- and adverse-risk groups. Cross-cohort analyses identified four bone marrow (BM) editing sites in *TMEM165, COQ4, TIMM17A*, and *PLXDC2* reproducibly associated with relapse and one peripheral blood (PB) editing site in *ABHD18* elevated in higher-risk ELN-2022 groups. Most editing sites were shared between BM and PB; only 2.1%-2.3% exhibited tissue-specific differences. Higher global editing levels were correlated with leukemic state, white blood cell count, and selected clinical features. These findings identify reproducible RNA editing signatures linked to AML disease stage and risk and support the use of RNA editing biomarkers for PB disease assessment.

## Introduction

Acute myeloid leukemia (AML) is a clinically heterogeneous hematologic malignancy characterized by diverse genetic and epigenetic alterations that drive variability in disease progression and treatment response^1^. Although large-scale genomic studies have substantially improved disease classification and risk stratification, relapse remains common and continues to be the major cause of treatment failure and mortality in AML^1^, highlighting the need to identify additional molecular mechanisms associated with disease progression and therapeutic resistance.

RNA editing, particularly adenosine-to-inosine (A-to-I) editing mediated by ADAR enzymes, is a pervasive post-transcriptional mechanism that can modulate RNA stability, splicing, coding potential, and immune signaling^2-4^. Emerging evidence suggests that RNA editing exerts context-dependent roles across cancers. Site-specific recoding can alter protein function (e.g., edited *AZIN1* has enhanced substrate binding affinity)^5^, editing can influence RNA splicing and miRNA biogenesis or targeting^6,7^, and widespread editing of endogenous double-stranded RNAs can modulate innate immune sensing^8,9^. Accordingly, ADAR activity has been associated with both oncogenic and tumor-suppressive effects and may influence responses to immunotherapies and epigenetic agents. Despite growing recognition of its biological importance, the contribution of RNA editing to AML pathogenesis and clinical heterogeneity remains incompletely understood. Unlike gene expression analyses, RNA editing is quantified at single-nucleotide resolution, requiring specialized analytical approaches for accurate site discovery and differential editing analysis.

Evidence supporting a role for RNA editing in AML remains limited. Early work demonstrated that RNA hyperediting at a pre-mRNA branch point in *PTPN6* synthesis disrupts splicing and produces an intron-retaining transcript enriched at diagnosis compared with remission^10^. More recently, large-scale analyses have cataloged thousands of editing events and reported associations between elevated global editing levels, adverse clinical outcomes, and specific genetic subtypes^11^. In addition, mechanistic studies have shown that core-binding factor fusions repress *ADARB1* transcription and that *ADARB1* can function as an editing-dependent tumor suppressor in specific AML contexts^12^. Together, these studies support a role for RNA editing in AML biology but leave several clinically important questions unresolved. Specifically, it remains unclear whether RNA editing signatures are associated with disease progression from diagnosis to relapse, whether they vary across ELN-2022 risk groups used for clinical risk stratification, and whether clinically informative editing signals can be reliably detected in peripheral blood. Addressing these questions is challenging because RNA editing is quantified at individual nucleotide positions and is highly dependent on accurate alignment, sequencing depth, local read coverage, and robust statistical modeling of count-based editing data.

To address these gaps, we analyzed RNA editing in the Epigenomics Studies in Acute Myeloid Leukemia (ESAML) cohort, which includes paired diagnosis and relapse samples together with age-matched healthy controls. Using a stringent RNA editing discovery pipeline and a beta-binomial regression framework that accounts for overdispersion and repeated measurements, we sought to identify RNA editing signatures associated with disease stage, including relapse, and ELN-2022 risk groups. We further evaluated the reproducibility of these signatures across independent AML cohorts and assessed the extent to which clinically relevant editing signals are shared between bone marrow (BM) and peripheral blood (PB). Through this approach, we provide a comprehensive evaluation of disease stage- and risk-associated RNA editing signatures in AML and examine their potential utility for PB-based disease assessment.

## Results

### Identification and Characterization of High-Confidence RNA Editing Sites in the ESAML Cohort

The ESAML cohort included paired diagnosis and relapse samples together with age-matched healthy controls, providing a framework to evaluate RNA editing across disease progression from normal hematopoiesis to AML diagnosis and relapse (**Supplementary Figure S1; Table 1**). After quality-control filtering, 126 samples remained for analysis, including 93 BM and 33 PB samples.

**Table 1.** Clinical and demographic characteristics of the study samples.

| Sample Stage | Total N | Bone Marrow |  |  |  | Peripheral Blood |  |  |  |
| --- | --- | --- | --- | --- | --- | --- | --- | --- | --- |
|  |  | BM N | Age Mean[min-max] | Sex |  | PB N | Age Mean[min-max] | Sex |  |
|  |  |  |  | F | M |  |  | F | M |
| **AML (Combined Diagnosis and Relapse)** | 118 | 85 | 50.5[18-74] | 34 | 51 | 33 | 53.0[25-74] | 8 | 25 |
| **Diagnosis** | 59 | 39 | 51.4[18-71] | 16 | 23 | 20 | 49.3[25-74] | 5 | 15 |
| **Relapse** | 59 | 46 | 49.8[20-74] | 18 | 28 | 13 | 58.7[42-74] | 3 | 10 |
| **Healthy Individuals** | 8 | 8 | 49.0[25-77] | 2 | 6 | — | — | — | — |

**Abbreviations:** BM, bone marrow; PB, peripheral blood; F, female; M, male.
**Notes:** — indicates not available. Light gray shading denotes BM samples, and dark gray shading denotes PB samples. N denotes the number of samples.

To address the technical challenges of RNA editing analysis from bulk RNA sequencing data, we applied a stringent RNA editing discovery framework incorporating dual-aligner validation, artifact filtering, and exclusion of known genomic variants (see **Supplementary Methods**). Candidate editing sites were identified using concordant alignments from STAR^13^ and Bowtie^14^ and subsequently filtered to remove sites overlapping known SNPs, repetitive regions, and other potential sequencing or mapping artifacts. Downstream analyses were restricted to recurrent sites detected in at least 5% of samples.

Using this framework, we identified 166,323 high-confidence RNA editing sites mapping to 5,917 genes. Most sites were located within intronic regions, followed by 3′ untranslated regions (3′UTRs), consistent with the known genomic distribution of A-to-I RNA editing^6,7,11,15^. All identified sites were cataloged in REDIportal v3.0^16^, supporting the robustness of the discovery framework and the high confidence of the detected editing events.

Functional enrichment analyses demonstrated that genes harboring intronic editing sites were enriched in cellular transport and metabolic pathways, whereas genes containing 3′UTR editing sites were enriched in stress-response and signaling pathways (**Supplementary Tables S2-S3**). This high-confidence RNA editing resource provided the foundation for subsequent analyses of disease stage-associated, risk-associated, and tissue-specific RNA editing signatures in AML.

### Disease Stage-Associated RNA Editing Signatures in AML

We next investigated whether RNA editing signatures were associated with AML disease stage, including healthy controls, diagnosis samples, and relapse samples. Using beta-binomial regression models with diagnosis as the reference group and adjusting for demographic, technical, and interferon-related covariates, we identified thousands of editing sites associated with disease stage (FDR < 0.1, a threshold widely used in differential editing analysis^17,18^; **Supplementary Table S4**), supporting dynamic changes in RNA editing during AML progression.

We further examined overlap among editing sites significantly associated with both healthy and relapse stages relative to diagnosis. Among these sites, 12.9% overlapped at the site level and 36.2% overlapped at the gene level (**Figure 1a-1b; Supplementary Table S5**). Notably, 107 overlapping sites exhibited opposite effect directions in the healthy-versus-diagnosis and relapse-versus-diagnosis comparisons (**Supplementary Table S6**). Specifically, sites with increased editing in healthy samples relative to diagnosis tended to show decreased editing in relapse relative to diagnosis, and vice versa. This pattern suggests substantial remodeling of RNA editing programs across disease progression.

**Figure 1.**
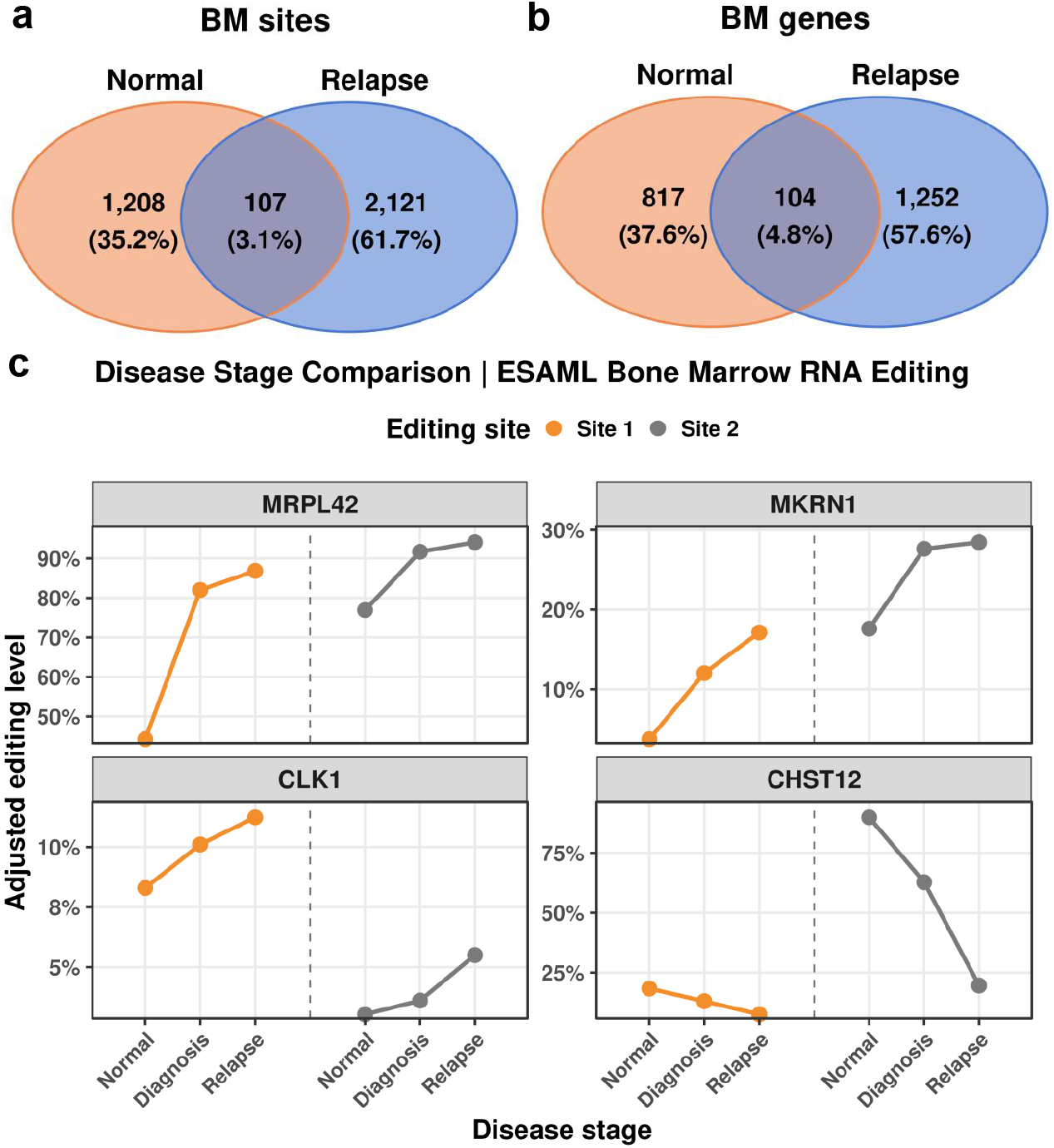
RNA editing is associated with disease stage. (a, b) Overlap of significantly associated RNA editing sites (a) and their host genes (b) between the Normal and Relapse groups in the ESAML bone marrow (BM) cohort. The Normal group represents healthy control individuals. (c) Distribution of eight editing sites from four genes across disease stages in ESAML BM samples. The y-axis shows the model-predicted mean editing level estimated from the fitted regression model. The eight sites include 7:2447915:+ (Site 1) and 7:2449394:+ (Site 2) in *CHST12*; 2:200855852:- (Site 1) and 2:200856402:- (Site 2) in *CLK1*; 7:140461511:- (Site 1) and 7:140464999:- (Site 2) in *MKRN1*; and 12:93472358:+ (Site 1) and 12:93474991:+ (Site 2) in *MRPL42*. Editing sites are denoted in the format of chromosome:position:strand.

Among these oppositely directed sites, 56 exhibited progressive increases in editing levels from healthy controls to diagnosis and subsequently to relapse. Notably, two sites each were located in *MRPL42, MKRN1*, and *CLK1*, genes involved in mitochondrial protein translation and cellular metabolism (*MRPL42*), ubiquitin-mediated protein regulation and RNA metabolism (*MKRN1*), and RNA splicing control (*CLK1*) (**Figure 1c**). In contrast, two editing sites within *CHST12* exhibited progressively decreased editing levels across disease stages. Together, these findings suggest that RNA editing signatures undergo coordinated remodeling during AML evolution and highlight candidate editing events that may contribute to disease progression and relapse.

### Relapse-Associated RNA Editing Signatures Are Reproducible Across Independent Cohorts

To evaluate the reproducibility of relapse-associated RNA editing signatures, we analyzed the independent GAML cohort (see Methods), which includes BM samples collected at diagnosis (day 0; d0), day 10 after treatment initiation (d10), and relapse (Rel). We focused on relapse-associated editing signals identified in ESAML and assessed their concordance across cohorts.

A total of 45 relapse-associated editing sites were shared between ESAML and GAML, and 24.2% of host genes harboring relapse-associated sites overlapped between cohorts (**Supplementary Table S7**). Among the 45 shared sites, 24 exhibited concordant effect directions, including 17 favoring relapse (**Supplementary Table S8**). Similar findings were observed when ESAML relapse samples were compared with GAML d10 samples, yielding 23.8% overlap at the gene level and 22 concordant sites, 19 of which favored d10. These observations support the reproducibility of a subset of relapse-associated editing signatures across independent AML cohorts despite differences in patient populations and clinical sampling designs.

Four editing sites were consistently identified across both validation analyses (**Table 2; Figure 2a-2d**). These sites were located within *TMEM165, COQ4, TIMM17A*, and *PLXDC2*, genes involved in Golgi homeostasis, mitochondrial metabolism, protein translocation, and angiogenesis-related pathways, respectively. Editing levels at these sites demonstrated consistent relapse-associated patterns across cohorts, supporting their potential as candidate biomarkers of disease progression and relapse.

**Table 2.**
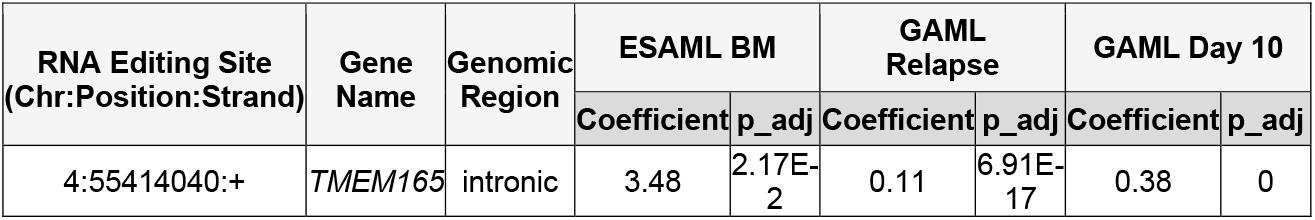
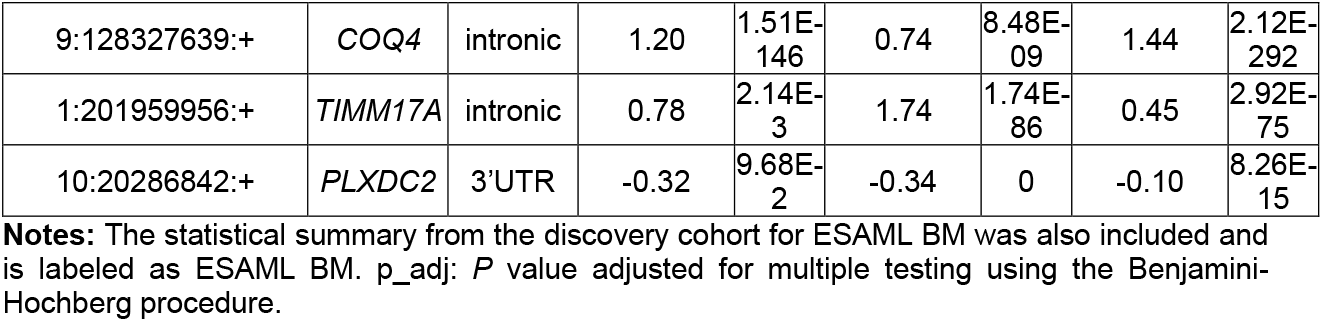
RNA editing sites validated in the GAML cohort, with adjusted *P* values < 0.05, and shared between the relapse and day 10.

| RNA Editing Site (Chr:Position:Strand) | Gene Name | Genomic Region | ESAML BM |  | GAML Relapse |  | GAML Day 10 |  |
| --- | --- | --- | --- | --- | --- | --- | --- | --- |
|  |  |  | Coefficient | p_adj | Coefficient | p_adj | Coefficient | p_adj |
| 4:55414040:+ | *TMEM165* | intronic | 3.48 | 2.17E-2 | 0.11 | 6.91E-17 | 0.38 | 0 |
| 9:128327639:+ | COQ4 | intronic | 1.20 | 1.51E-146 | 0.74 | 8.48E-09 | 1.44 | 2.12E-292 |
| 1:201959956:+ | TIMM17A | intronic | 0.78 | 2.14E-3 | 1.74 | 1.74E-86 | 0.45 | 2.92E-75 |
| 10:20286842:+ | PLXDC2 | 3'UTR | -0.32 | 9.68E-2 | -0.34 | 0 | -0.10 | 8.26E-15 |

**Notes:** The statistical summary from the discovery cohort for ESAML BM was also included and is labeled as ESAML BM. p\_adj: *P* value adjusted for multiple testing using the Benjamini-Hochberg procedure.

**Figure 2.**
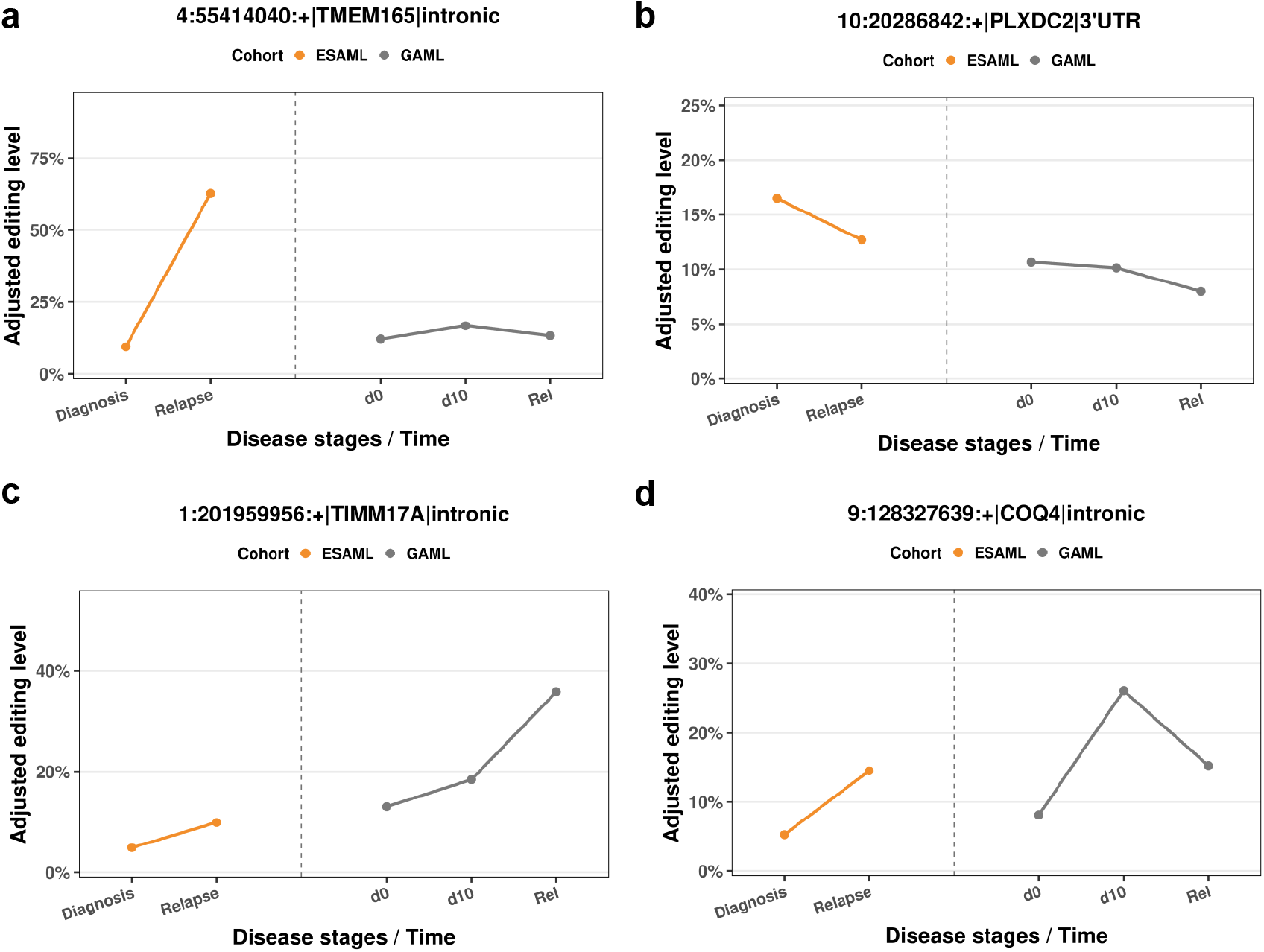
Four examples of RNA editing sites associated with relapse. Distribution of four editing sites from four genes across disease stages in ESAML bone marrow (BM) samples (left) and post-treatment time points in GAML (right), including d0 (diagnosis), d10 (day 10 post-treatment), and Rel (relapse). Y-axis is the predicted mean editing level for each site, estimated from the fitted regression model. The four sites include 1:201959956:+ in *TIMM17A*, 4:55414040:+ in *TMEM165*, 9:128327639:+ in *COQ4*, and 10:20286842:+ in *PLXDC2*. Editing sites are denoted in the format of chromosome:position:strand.

### RNA Editing Signatures Associated with ELN-2022 Risk Stratification

We next investigated whether RNA editing signatures were associated with ELN-2022 risk stratification, the current standard for clinical risk assessment in AML. Across both BM and PB samples, hundreds to thousands of editing sites showed significant associations with ELN-2022 classification (FDR < 0.1; **Supplementary Table S4**), indicating that RNA editing patterns vary across clinically relevant AML risk categories.

Among ELN-associated editing signatures, the greatest overlap was observed between the Intermediate- and Adverse-risk groups. At the site level, 23.4% of significant sites overlapped in PB and 18.5% in BM, whereas gene-level overlap reached 44.6% in PB and 39.6% in BM (**Supplementary Table S5**). Most overlapping sites exhibited concordant effect directions, with more than 76.1% showing associations in the same direction across both risk groups (**Figure 3a-3d**). These findings suggest that Intermediate- and Adverse-risk AML share substantial overlapping RNA editing programs.

**Figure 3.**
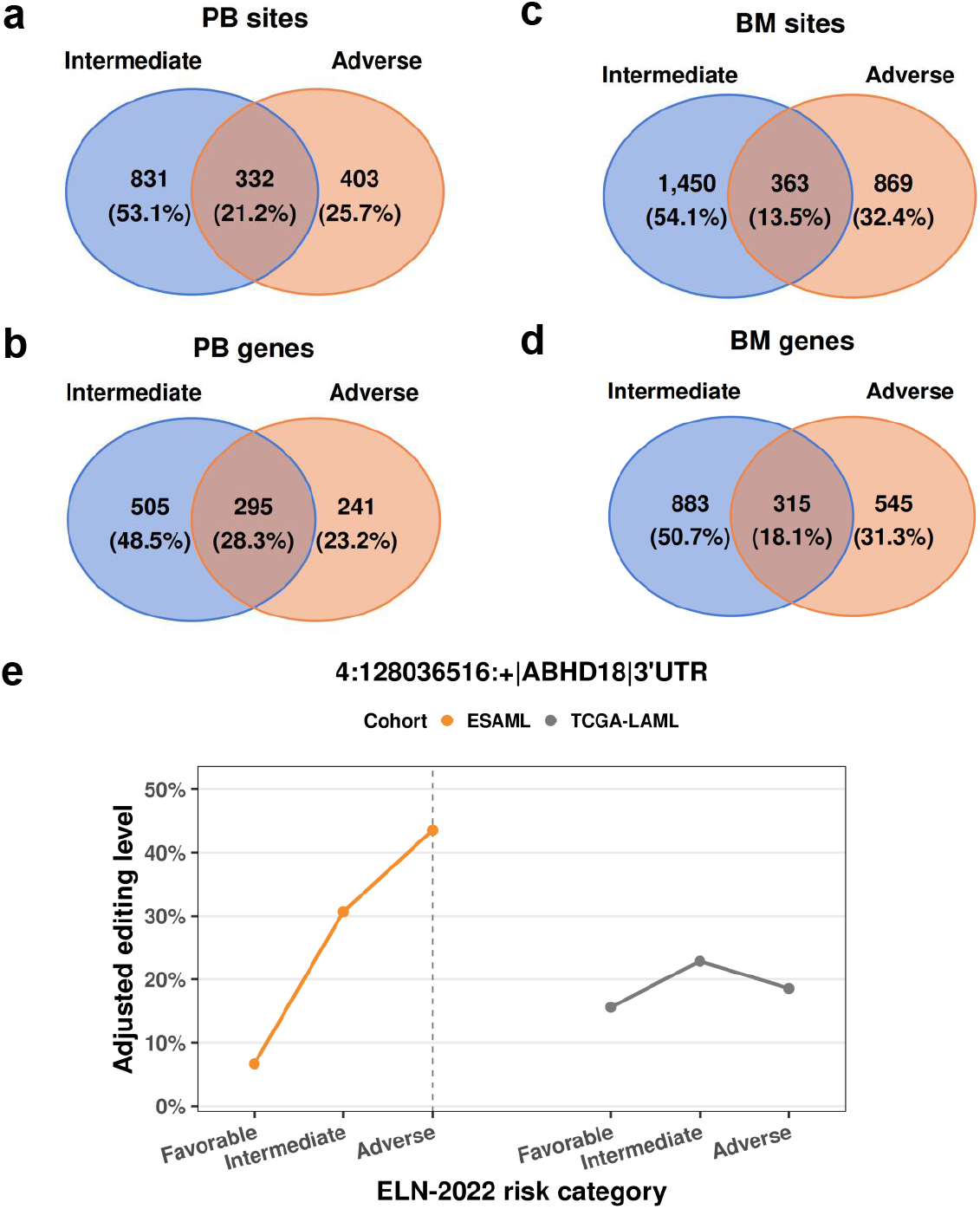
RNA editing is associated with ELN-2022 risk category. (a-d) Overlap of significantly associated editing sites (a, c) and their host genes (b, d) between the ELN-2022 Intermediate and Adverse groups in peripheral blood (PB; a, b) and bone marrow (BM; c, d). (e) Distribution of the editing site (chr4:128036516:+) across ELN-2022 categories in ESAML PB samples (left) and TCGA samples (right). Y-axis is the predicted mean editing level for each site, estimated from the fitted regression model.

To evaluate reproducibility, we compared ELN-associated editing signatures identified in ESAML PB samples with those observed in the independent TCGA-LAML cohort. Although overlap at the individual-site level was limited, 14.5% of host genes harboring significant ELN-associated editing sites were shared between cohorts (**Supplementary Table S9**), indicating moderate gene-level concordance despite differences in cohort composition and sequencing protocols. These findings further highlight the challenges of reproducing site-specific RNA editing signals across independent studies while supporting the robustness of gene-level editing programs associated with AML risk stratification.

Among the shared signals, a 3′UTR editing site in *ABHD18* (chr4:128036516) was significantly associated with both Intermediate- and Adverse-risk groups in ESAML and TCGA-LAML (**Table 3; Figure 3e**). Increased editing at this site was consistently associated with higher-risk categories across cohorts. Notably, this association was detected in PB but not BM, highlighting the potential utility of selected RNA editing events as PB-accessible biomarkers for AML risk stratification.

**Table 3.** RNA editing sites validated in the TCGA-LAML cohort, with adjusted *P* values < 0.05, and shared between the ELN-2022 intermediate- and adverse-risk groups.

| RNA Editing Site (Chr:Position:Strand) | Gene Name | Genomic Region | ELN-2022 Risk Groups | Coefficient | SE | p_adj | Cohorts |
| --- | --- | --- | --- | --- | --- | --- | --- |
| 4:128036516:+ | ABHD18 | 3'UTR | Intermediate | 0.48 | 0.07 | 2.70E-10 | TCGA-LAML |
| 4:128036516:+ | ABHD18 | 3'UTR | Intermediate | 3.29 | 0.47 | 7.06E-11 | ESAML_PB |
| 4:128036516:+ | ABHD18 | 3'UTR | Adverse | 0.21 | 0.01 | 4.24E-88 | TCGA-LAML |
| 4:128036516:+ | ABHD18 | 3'UTR | Adverse | 4.33 | 0.45 | 3.62E-20 | ESAML_PB |

**Abbreviations:** SE, standard error. Chr, chromosome.
**Notes:** The statistical summary from the discovery cohort for ESAML PB was also included and is labeled as ESAML PB. p\_adj: *P* value adjusted for multiple testing using the Benjamini-Hochberg procedure.

### RNA Editing Landscapes Are Largely Conserved Between Bone Marrow and Peripheral Blood

Because BM sampling is invasive and may not be feasible for serial disease monitoring, we next evaluated whether PB captures RNA editing patterns observed in BM. We therefore compared RNA editing landscapes across tissues and assessed the extent of shared editing signals.

Most RNA editing sites were shared between BM and PB. Across diagnosis and relapse samples in the ESAML cohort, 77.5% of editing sites were detected in both tissues, corresponding to 94.6% overlap at the gene level (**Figure 4a-b; Supplementary Table S10**). Similar patterns were observed in the independent BEAT-AML cohort, where 85.7% of editing sites and 97.5% of host genes were shared between BM and PB samples. In both ESAML and BEAT-AML, RNA editing sites exhibited highly similar genomic distributions across BM and PB samples, with the majority of sites located within intronic regions followed by 3’ untranslated regions (3′UTRs) (**Figure 4c-d**). These findings demonstrate substantial concordance of RNA editing landscapes across tissues and support the reproducibility of BM-PB overlap across independent AML cohorts.

**Figure 4.**
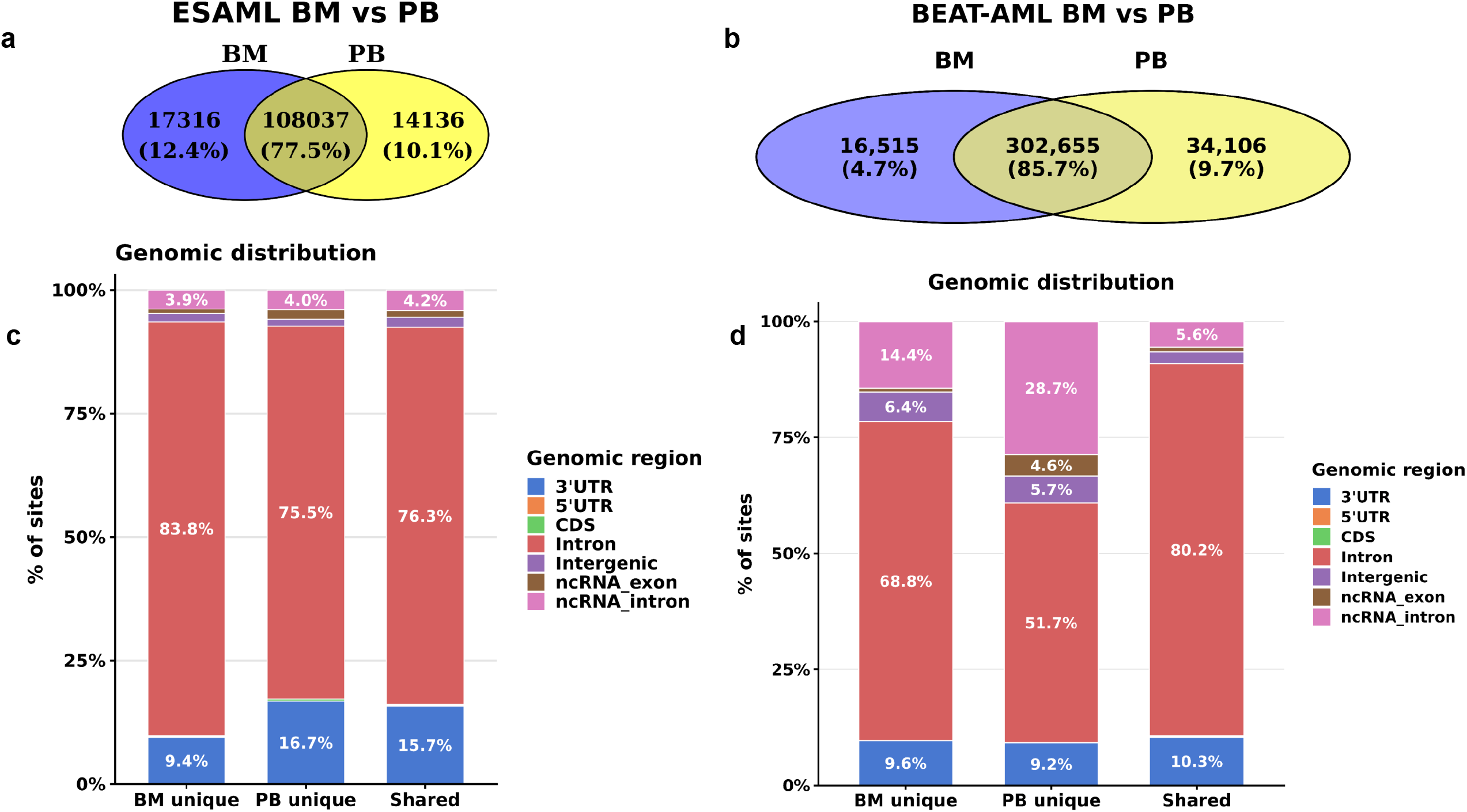
Comparison of RNA editing sites between bone marrow and peripheral blood in ESAML and BEAT-AML. (a, b) Overlap of RNA editing sites identified in bone marrow (BM) and peripheral blood (PB) across all AML samples in ESAML (a) and BEAT-AML (b). (c, d) Genomic distribution of site categories in ESAML (c) and BEAT-AML (d). Only tumor samples were included in these analyses; healthy control samples were excluded. BM-unique and PB-unique sites in (c) and (d) correspond to the unique site sets shown in (a) and (b), respectively; shared sites correspond to the overlap between BM and PB.

To identify tissue-associated editing events, we performed differential editing analyses between BM and PB samples while accounting for repeated measurements and relevant clinical covariates. Only 2.1%–2.3% of evaluated sites exhibited significant tissue-associated differences (FDR < 0.1; **Supplementary Table S10**), indicating that the vast majority of RNA editing events are conserved between BM and PB. Although tissue-associated differences were more evident at the individual-site level than at the gene level, overall RNA editing patterns remained highly similar across tissues.

We next examined whether clinically associated RNA editing signatures were shared between BM and PB. Across disease stage and ELN-2022 analyses, overlap of significant editing sites was limited at the individual-site level (0.6%–0.8%) but substantially greater at the gene level (21.0%–28.7%) (**Supplementary Table S11**). Notably, these overlap rates were comparable to those observed across clinical variables and independent cohorts throughout the study, suggesting that differences between BM and PB are not substantially greater than other sources of biological and technical variability affecting site-level RNA editing analyses.

Importantly, the substantial overlap between BM and PB editing landscapes indicates that PB captures a large proportion of the RNA editing repertoire present in BM. In addition, PB analyses identified clinically associated editing events, including an ELN-associated editing site within *ABHD18*, demonstrating that PB-derived RNA editing measurements can provide clinically informative biomarkers regardless of whether individual editing sites are shared with BM. Together, these findings support the potential utility of PB as a minimally invasive source for RNA editing-based biomarker discovery, disease assessment, and longitudinal monitoring in AML.

### Global RNA Editing Burden Is Associated with Disease Burden and Selected Clinical Features

To determine whether the site-level associations identified above were accompanied by broader changes in RNA editing activity, we evaluated global RNA editing levels across clinical variables. Global editing levels were calculated as the mean editing proportion across all evaluated sites within each sample and tested for association with clinical features.

In ESAML, higher global editing levels in BM were positively associated with white blood cell (WBC) count and negatively associated with the normal disease stage (**Supplementary Figure 2a-2b; Supplementary Table S12**), indicating increased RNA editing activity in leukemic BM samples, particularly those with greater disease burden.

These findings were supported by independent AML cohorts. In BEAT-AML, higher BM global editing levels were associated with adverse ELN-2022 risk and refractory disease (**Supplementary Figure 2c; Supplementary Table S12**). In GAML, global editing levels were significantly lower in normal BM samples than in AML samples (**Supplementary Figure 2d; Supplementary Table S12**).

Together, these findings suggest that increased global RNA editing activity is associated with leukemic state, disease burden, and selected adverse clinical features. In contrast to the site-level analyses, associations with ELN-2022 risk were less consistently observed, indicating that clinically relevant RNA editing differences are more readily captured at the individual-site level than by global editing burden alone.

## Discussion

In this study, we systematically characterized RNA editing signatures associated with AML disease stage, relapse, and ELN-2022 risk stratification using a stringent RNA editing discovery framework and count-based beta-binomial regression modeling. We identified thousands of RNA editing sites associated with clinically relevant disease characteristics and validated a subset of relapse- and risk-associated signals in independent cohorts. Although site-level RNA editing analyses are inherently challenging because of technical and biological variability, several editing events demonstrated reproducible associations across datasets. We also observed substantial concordance of RNA editing landscapes between BM and PB, indicating that many editing signals can be detected outside the primary disease site. Collectively, these findings establish RNA editing as a clinically relevant dimension of molecular heterogeneity in AML and support its potential utility for biomarker development and disease assessment.

A major finding of this study was the identification of RNA editing programs associated with AML disease progression. While previous studies have linked global RNA editing activity to AML outcome and genetic subtypes, our analyses demonstrate that disease-stage associations are also evident at the individual-site level and can be detected across the continuum from healthy hematopoiesis to diagnosis and relapse. Notably, multiple editing sites exhibited progressive increases or decreases across disease stages, suggesting that RNA editing programs undergo dynamic remodeling during AML evolution rather than representing static disease-associated features. These observations are consistent with growing evidence that post-transcriptional regulatory mechanisms contribute to leukemic adaptation and disease progression and suggest that RNA editing may provide biological insights complementary to those obtained from genomic and transcriptomic biomarkers.

Another important finding was the identification of relapse- and risk-associated editing signatures that were reproducible across independent AML cohorts. Reproducibility at the individual-site level remains a significant challenge in RNA editing studies because editing measurements are highly sensitive to alignment accuracy, sequencing depth, local read coverage, and biological heterogeneity. Despite these challenges, we identified relapse-associated editing sites within *TMEM165, COQ4, TIMM17A*, and *PLXDC2* that consistently distinguished relapsed disease across cohorts, as well as an ELN-associated editing site within *ABHD18* that replicated in both ESAML and TCGA-LAML. Although the functional consequences of these editing events remain unclear, their recurrence across independent datasets suggests that they represent biologically meaningful features rather than cohort-specific observations and highlights the potential value of RNA editing as a source of clinically informative biomarkers.

An important translational finding of this study was the extensive conservation of RNA editing landscapes between BM and PB. Although BM remains the primary tissue for AML diagnosis and monitoring, repeated BM sampling is invasive and may not always be feasible, particularly in longitudinal studies and clinical surveillance settings. We found that most editing sites and edited genes were shared between BM and PB, whereas only a small fraction of sites exhibited tissue-associated differences. These findings indicate that a substantial proportion of AML-associated RNA editing activity is preserved across tissues and that PB captures much of the RNA editing repertoire present in BM. Importantly, the clinical utility of PB-derived RNA editing biomarkers does not depend entirely on concordance with BM. Editing events detected exclusively or preferentially in PB may still provide clinically informative biomarkers even if they do not directly reflect the underlying biology of BM. Thus, the value of PB-based RNA editing assessment may arise from both shared editing programs that may mirror disease-associated processes in BM and PB-specific editing signatures that contribute independent clinical information. Together, these observations support further investigation of RNA editing–based approaches for minimally invasive disease assessment and biomarker development in AML.

Our findings also underscore both the opportunities and analytical challenges associated with RNA editing studies in AML. Unlike gene expression measurements, RNA editing is quantified at individual nucleotide positions and is therefore particularly sensitive to sequencing depth, local coverage, alignment accuracy, and biological heterogeneity. To mitigate these challenges, we employed a stringent discovery framework incorporating dual-aligner validation, extensive artifact filtering, exclusion of known genomic variants, and beta-binomial regression models that account for overdispersion and repeated measurements. This strategy enabled the identification of high-confidence RNA editing events while reducing false-positive discoveries. The successful replication of several relapse- and risk-associated editing signatures across independent cohorts further supports the utility of rigorous site-level analytical approaches for RNA editing biomarker discovery.

An additional observation from this study is that overlap across tissues and independent cohorts was generally greater at the gene level than at the individual-site level. This pattern likely reflects both technical variability in site-level quantification and biological heterogeneity in RNA editing regulation. Although beta-binomial regression effectively models count-based editing measurements, the stronger concordance observed at the gene level suggests that future studies may benefit from integrating information across multiple editing sites within the same gene, transcript, or regulatory region rather than relying exclusively on individual editing events. Such approaches may improve robustness while preserving biological interpretability.

Several limitations should be considered. First, although this study represents one of the largest evaluations of RNA editing in AML to date, the number of matched diagnosis-relapse samples remains limited, reflecting the scarcity of longitudinal AML cohorts with high-quality RNA sequencing data. Second, the biological functions of most identified editing events remain unknown and will require mechanistic investigation. Third, because our analyses were performed using bulk RNA sequencing data, the observed editing signals likely reflect a combination of leukemic and non-leukemic cell populations. Future studies incorporating single-cell and multi-omic approaches will be important for resolving cell type-specific editing programs and determining their functional consequences during AML progression.

In summary, we demonstrate that RNA editing landscapes are extensively remodeled during AML progression, identify reproducible editing signatures associated with relapse and ELN-2022 risk, and show that many RNA editing signals are conserved between BM and PB. These findings establish RNA editing as a promising source of AML biomarkers and provide a foundation for future studies aimed at elucidating the biological functions, clinical utility, and therapeutic implications of RNA editing-based signatures in AML.

## Methods

### Study Cohorts

The Epigenomics Studies in Acute Myeloid Leukemia (ESAML) cohort (dbGaP accession: phs001027) served as the primary discovery cohort. Independent validation analyses were performed using three additional AML cohorts: The Cancer Genome Atlas Acute Myeloid Leukemia (TCGA-LAML; dbGaP accession: phs000178), the Beat Acute Myeloid Leukemia Master Trial (BEAT-AML; dbGaP accession: phs001657), and the Genomics of Acute Myeloid Leukemia (GAML; dbGaP accession: phs000159). Clinical and demographic information for all cohorts was obtained from the corresponding study repositories.

### RNA Editing Site Discovery and Quantification

RNA editing site-level read counts were generated using a comprehensive RNA editing discovery pipeline incorporating dual-alignment validation and artifact filtering. Briefly, sequencing reads were aligned to the GRCh38 reference genome using complementary alignment strategies based on STAR^13^ and Bowtie^14^. Candidate editing sites were required to have concordant alignments across both methods and were subsequently filtered to remove sites overlapping known genomic variants, repetitive regions, and other potential sequencing or mapping artifacts. Additional filtering was performed to exclude sites exhibiting significant strand bias or positional bias within sequencing reads. To further enrich for high-confidence RNA editing events, sites exhibiting saturated editing patterns, defined as an editing ratio of 1 in more than 95% of samples, were excluded. Downstream analyses were restricted to recurrent editing sites detected in at least 5% of samples.

RNA editing levels were quantified as the proportion of edited reads among total reads covering each site. Edited-read counts and total read counts were retained for downstream statistical analyses.

### Differential RNA Editing Analysis

Because RNA editing is measured as edited-read counts relative to total read coverage and frequently exhibits greater variability than expected under a simple binomial distribution, associations between RNA editing and clinical variables were evaluated using beta-binomial regression models implemented in the glmmTMB package^19^. These models account for overdispersion while preserving read-count information.

To account for interferon-related effects on RNA editing, we curated a panel of 127 interferon-related and interferon-stimulated genes from our another study^20^ and derived the first two principal components (PC1_ISG and PC2_ISG). Age, sex, library selection type, PC1_ISG, and PC2_ISG were included as covariates in all association models.

Site-specific associations with ELN-2022 risk stratification, disease stage, white blood cell (WBC) count, and BM-versus-PB tissue source were evaluated using beta-binomial regression. Favorable risk was used as the reference level for ELN-2022 analyses, and diagnosis was used as the reference level for disease-stage analyses. When repeated measurements were available, subject-level random effects were included to account for within-subject correlation. P values were adjusted across sites using the Benjamini-Hochberg method, and sites with a false discovery rate (FDR) < 0.10 were considered statistically significant, a commonly used threshold^17,18^.

### Global RNA Editing Analysis

Global RNA editing burden was defined as the ratio of summed edited reads to summed total reads across all retained editing sites within each sample. Associations between global editing burden and clinical variables were evaluated using beta-binomial models adjusted for the same covariates described above.

### Functional Enrichment and Cross-Cohort Validation

Functional enrichment analyses were performed for genes harboring significant RNA editing sites using gene ontology and pathway-based approaches. Cross-cohort validation analyses were conducted by comparing significant editing sites and host genes identified in the ESAML cohort with corresponding results from the TCGA-LAML, GAML, and BEAT-AML cohorts. The same RNA editing discovery pipeline and statistical association framework were applied across all cohorts.

Additional details regarding data preprocessing, filtering criteria, statistical models, and software versions are provided in the **Supplementary Methods**.

## Acknowledgements

We thank Emily Diaz, PhD, for excellent assistance with the preparation of the manuscript. This work is supported, in part, by Institutional Research Grant IRG #22-151-37-IRG from the American Cancer Society and by the Medical College of Wionsin (MCW) Cancer Center, and Versiti Blood Research Institute Foundation.

