## Supplementary figures and images for "Disease Stage- and Risk-Associated RNA Editing Signatures in Acute Myeloid Leukemia and Their Utility for Peripheral Blood-Based Assessment"

### Supplementary Figures1-2

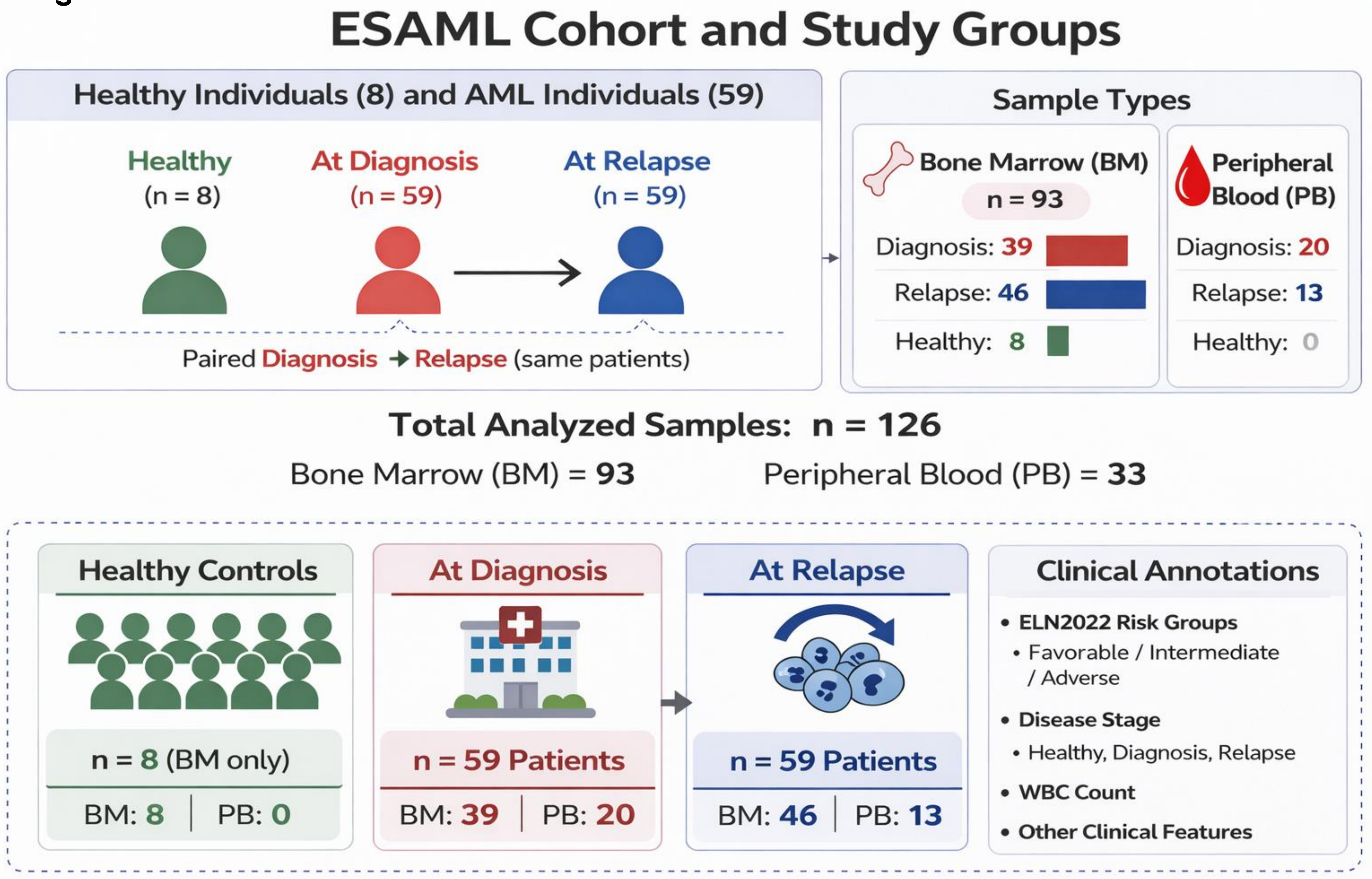

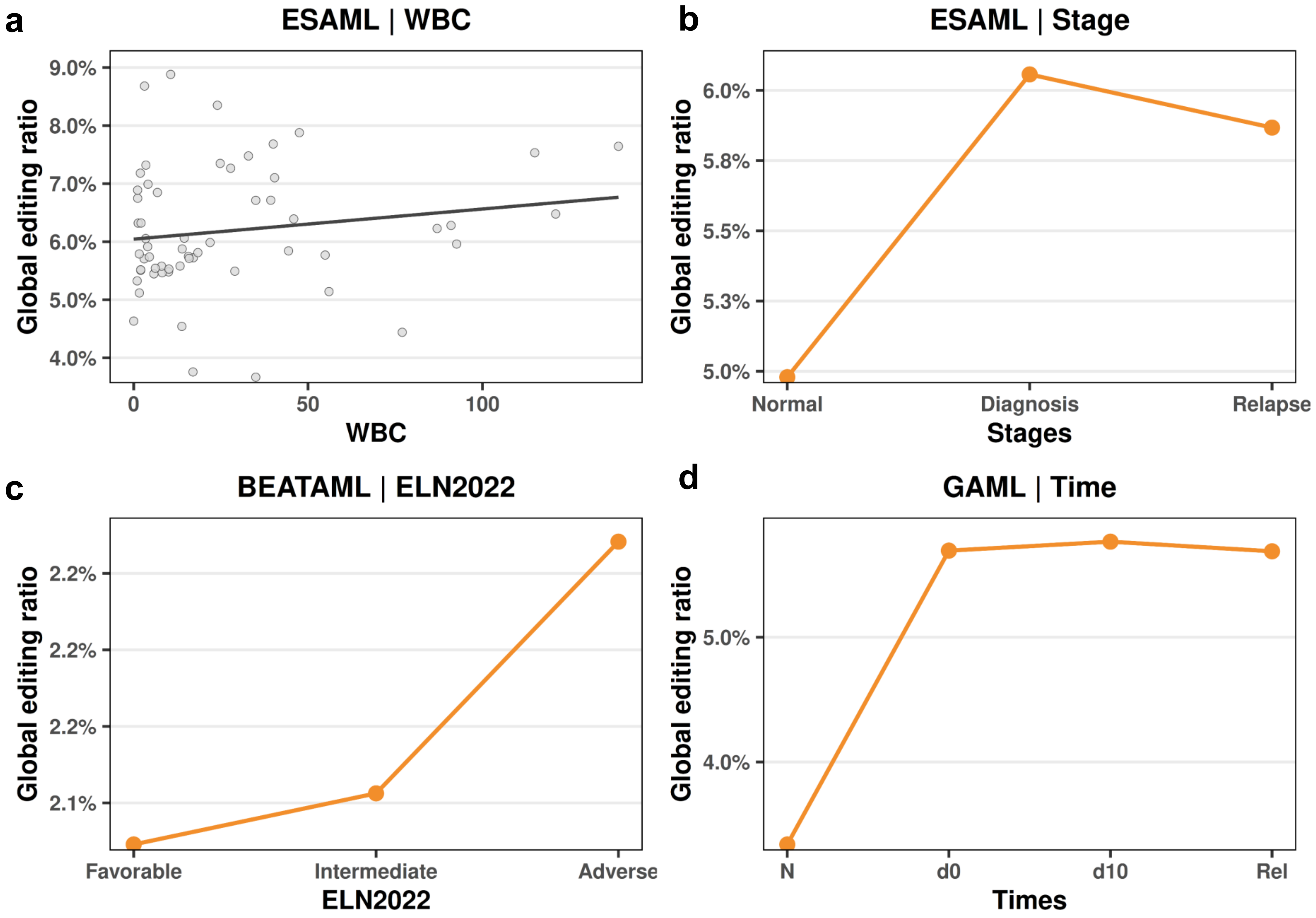
