## Supplementary Methods for "Disease Stage- and Risk-Associated RNA Editing Signatures in Acute Myeloid Leukemia and Their Utility for Peripheral Blood-Based Assessment"

### ESAML cohort and RNA editing data processing

Raw RNA sequencing data and corresponding clinical and demographic information were obtained from the Epigenomics Studies in Acute Myeloid Leukemia (ESAML) cohort (dbGaP accession: phs001027). To enable robust site-level RNA editing analyses, we developed a comprehensive RNA editing discovery and quantification pipeline designed to minimize technical artifacts while preserving sensitivity for detecting bona fide A-to-I editing events. The pipeline integrates complementary alignment strategies, multiple layers of artifact filtering, and count-based quantification to generate high-confidence editing measurements suitable for downstream statistical modeling.

Briefly, sequencing reads were quality assessed and aligned to the GRCh38 reference genome using complementary STAR[1] two-pass and Bowtie[2] alignment strategies. Candidate editing sites were required to be supported by concordant alignments across both methods, reducing potential mapping-related artifacts. Sites were subsequently filtered to remove loci overlapping known genomic variants, repetitive regions, ribosomal RNA annotations, and other sources of spurious variation. Additional quality-control procedures excluded sites exhibiting significant strand bias, positional bias, or nearby non–A-to-G mismatches, thereby enriching for high-confidence A-to-I RNA editing events. The final output consisted of a site-level matrix containing, for each editing site in each sample, the number of edited reads, the total number of reads, and the corresponding editing ratio.

To define a high-confidence set of editing sites for downstream analyses, additional cohort-level filters were applied. Sites were retained if editing was detected in at least 5% of samples and were excluded if more than 95% of informative samples exhibited an editing ratio of 1. These filters were intended to reduce the influence of low-frequency artifacts and sites likely to reflect underlying genomic variants rather than true RNA editing events. The retained sites were used for both site-level and global RNA editing analyses.

For each retained site, matrices of edited read counts, total read counts, and editing ratios were generated following harmonization of sample identifiers across molecular and clinical datasets. These count-based measurements served as the primary input for beta-binomial regression analyses of disease stage, relapse status, ELN-2022 risk groups, and bone marrow (BM)-peripheral blood (PB) comparisons.

### Clinical and gene expression data integration

Clinical annotations curated for the ESAML cohort included ELN-2022 risk category (Favorable, Intermediate, and Adverse), disease stage (Normal, Diagnosis, and Relapse), white blood cell count (WBC), age, and sex. Library selection type was included as a technical covariate to account for potential batch or protocol-related effects. Samples were included in a given analysis only if the relevant clinical variable, covariates, and editing measurements were all available.

Because ADAR1 p150, a major enzyme mediating A-to-I RNA editing, is inducible by interferon signaling[3], its expression may reflect inflammatory or interferon-driven transcriptional states in addition to RNA editing activity. To account for interferon-related transcriptional effects, interferon-stimulated gene (ISG) expression was summarized using principal component analysis (PCA) following our previously described approach[4]. Briefly, normalized expression values for a curated interferon-stimulated gene set (ISG) were scaled, and principal components were derived across samples. The first two components, PC1_ISG and PC2_ISG, were included as covariates in all clinical association models.

**Site-level association analysis with ELN-2022 risk category**

To assess whether site-specific RNA editing was associated with ELN-2022 risk category (ELN-2022), we modeled editing read counts at each site using a beta-binomial regression framework implemented in glmmTMB package[5], with a binomial model used as a fallback when the beta-binomial model failed to converge. For sample *i* at site *s*, let $E_{is}$ denote the number of edited reads and $T_{is}$ the total number of reads covering the site. The number of unedited reads was defined as

$$U_{is} = T_{is} - E_{is}.$$

For each site, we assumed:

$$E_{is} \sim \text{Binomial}(T_{is}, p_{is}),$$

where the latent probability $p_{is}$follows a Beta distribution, $p_{is} \sim \text{Beta}(a_{is}, b_{is})$.

Using the mean-precision parameterization,

$$a_{is} = \mu_{is}\phi_s, b_{is} = (1 - \mu_{is})\phi_s,$$

where $\mu_{is}$is the conditional mean editing proportion and $\phi_s$ is a site-specific precision parameter. Under this parameterization,

$$E\big(p_{is} \mid X_i, u_{\text{subject}(i)}\big) = \mu_{is}, \text{Var}(p_{is} | X_i, u_{\text{subject}(i)}) = \frac{\mu_{is}(1-\mu_{is})}{\phi_s+1}.$$

Marginally, this yields a beta-binomial distribution for the edited read count:

$$E_{is} \mid X_i, b_{\text{subject}(i)} \sim \text{Beta-binomial}(T_{is}, \mu_{is}, \phi_s),$$

with

$$E(E_{is} \mid X_i, b_{\text{subject}(i)}) = T_{is}\mu_{is}$$

and

$$\text{Var}(E_{is} \mid X_i, b_{\text{subject}(i)}) = T_{is}\mu_{is}(1 - \mu_{is})\frac{T_{is} + \phi_s}{\phi_s + 1}.$$

We modeled the mean editing proportion $\mu_{is}$using a logit link:

$$\text{logit}(\mu_{is}) = \beta_0 + \beta_1\text{ELN2022}_{\text{Intermediate},i} + \beta_2\text{ELN2022}_{\text{Adverse},i} + \beta_3\text{PC1\_ISG}_i + \beta_4\text{PC2\_ISG}_i + \beta_5\text{AGE}_i + \beta_6\text{SEX}_i + \beta_7\text{LibrarySelection}_i + b_{\text{subject}(i)},$$

where $b_{\text{subject}(i)} \sim N(0, \sigma_s^2)$is a subject-level random intercept included to account for within-subject dependence among multiple samples from the same individual. The precision parameter

$\phi_s$ was modeled as constant across samples for a given site. ELN-2022 was treated as a categorical predictor with Favorable risk as the reference group; thus, the resulting coefficients for ELN-2022 estimate differences in the mean editing proportion for Intermediate versus Favorable and Adverse versus Favorable risk groups, conditional on covariates. Only samples with nonmissing ELN-2022 annotation and informative coverage at the site were included. Sites with insufficient informative data after filtering were excluded from modeling.

For each fitted model, we extracted the coefficients and P values for the ELN-2022 contrast terms. P values were adjusted across sites using the Benjamini-Hochberg false discovery rate procedure. Sites with q < 0.10 for either ELN-2022 contrast were defined as ELN-associated editing sites.

**Site-level association analysis with disease stage and WBC**

A similar modeling framework was used to evaluate associations between site-specific RNA editing and disease stage or WBC. Disease stage was modeled as a categorical predictor, using Diagnosis as the reference category, whereas WBC was modeled as a continuous predictor.

**Quantification of global RNA editing burden**

To quantify sample-level global RNA editing burden, we aggregated read counts across all retained editing sites. For each sample *i*, the total number of edited reads was defined as the sum of edited reads across sites, and the total read count was defined as the sum of all reads across sites. The number of unedited reads was calculated as the difference between total and edited reads:

$$global\_edited_i = \sum_s E_{is}.$$
$$global\_total_i = \sum_s T_{is}.$$
$$global\_unedited_i = global\_total_i - global\_edited_i.$$

The global editing ratio was then calculated as:

$$global\_ratio_i = \frac{global\_edited_i}{global\_total_i}, \text{for samples with } global\_total_i > 0.$$

Only samples with $global\_total_i > 0$ were included in global editing analyses.

**Association of global RNA editing burden with ELN-2022 risk category, disease stage, and WBC**

We next evaluated whether global RNA editing burden was associated with ELN-2022 risk category, disease stage, and WBC. For ELN-2022 analyses, samples with nonmissing ELN-2022 annotation were modeled using a beta-binomial regression framework implemented in glmmTMB package:

For sample $i$, let $G_i$ denote the summed edited reads across all retained sites and $N_i$ the summed total reads across those sites, with $U_i = N_i - G_i$ denoting the summed unedited reads. Global RNA editing burden was modeled as

$$G_i \sim \text{Beta-binomial}(N_i, \mu_i, \phi),$$

with

$$\text{logit}(\mu_i) = \beta_0 + \beta_1 \text{ELN2022}_{\text{Intermediate},i} + \beta_2 \text{ELN2022}_{\text{Favorable},i} + \beta_3 \text{PC1}_{\text{ISG}_i} + \beta_4 \text{PC2}_{\text{ISG}_i} + \beta_5 \text{AGE}_i + \beta_6 \text{SEX}_i + \beta_7 \text{LibrarySelection}_i + b_{\text{subject}(i)}.$$

Here, $\mu_i$ represents the expected global editing proportion for sample $i$, and $\phi$ is the dispersion parameter, which captures extra-binomial variation beyond that expected under a binomial model. $b_{\text{subject}(i)}$ denotes a subject-specific random intercept included to account for repeated measures from the same individual. Under this parameterization, the underlying global editing probability $p_i$ is assumed to follow a Beta distribution:

$$p_i \sim \text{Beta}(a_i, b_i),$$

where

$$a_i = \mu_i \phi, b_i = (1 - \mu_i)\phi.$$

ELN-2022 was modeled as a categorical predictor with Adverse risk as the reference group. Accordingly, the resulting coefficients for the Favorable and Intermediate groups represent adjusted differences in the mean global RNA editing proportion relative to the Adverse group. PC1_ISG and PC2_ISG represent the first two principal components derived from the curated interferon-stimulated gene set.

A similar beta-binomial modeling framework was used for disease stage and WBC. Disease stage was modeled as a categorical predictor with Diagnosis as the reference category, whereas WBC was modeled as a continuous predictor. All models were adjusted for PC1_ISG, PC2_ISG, age, sex, and library selection type.

**Cohort description**

Raw RNA sequencing data and associated clinical and demographic information were obtained and analyzed from four independent AML cohorts: The Cancer Genome Atlas Acute Myeloid Leukemia (TCGA-LAML) (dbGaP accession: phs000178), the Beat Acute Myeloid Leukemia Master Trial (BEAT-AML) (dbGaP accession: phs001657), the Epigenomics Studies in Acute Myeloid Leukemia (ESAML) (dbGaP accession: phs001027), and the Genomics of Acute Myeloid Leukemia (GAML) (dbGaP: phs000159).
